# Nutrition Outweighs Defense: *Myzus Persicae* (Green Peach Aphid) Prefers and Performs Better on Young Leaves of Cabbage

**DOI:** 10.1101/085159

**Authors:** He-He Cao, Zhan-Feng Zhanga, Xiao-Feng Wang, Tong-Xian Liu

## Abstract

Plant leaves of different ages differ in nutrition and toxic metabolites and thus exhibit various resistance levels against insect herbivores. However, little is known about the relationship between leaf ontogeny and plant resistance against phloem-feeding insects. In this study, we found that the green peach aphid, *Myzus persicae* Sulzer, preferred to settle on young cabbage leaves (*Brassica oleracea* L. var. *capitata*) rather than mature or old leaves, although young leaves contained the highest concentration of glucosinolates. Furthermore, aphids feeding on young leaves had higher levels of glucosinolates in their bodies, but aphids performed better on young leaves in terms of body weight and population growth. The concentration of glutamine in young leaves was the highest, which stimulated aphids feeding when added to the sucrose solution. Phloem sap of young leaves had higher amino acid:sucrose molar ratio than mature leaves, and aphids feeding on young leaves showed two times longer phloem feeding time and five times more dry honeydew excretion than on other leaves. These results indicate that aphids acquired the highest amount of nutrition and defensive metabolites when feeding on young cabbage leaves that are strong natural plant sinks. The higher phloem sap availability of young leaves likely contributes to the attractiveness and suitability for aphids and may compensate the negative effects of glucosinolates on aphids. According to these findings, we propose that phloem sap availability influenced by leaf ontogeny and source-sink status play a significant role in plant-aphid interaction, which desires more attention in future research.

## INTRODUCTION

The optimal-defense hypothesis suggests that valuable tissues of a plant should be better defended against insect herbivores (Barton and Koricheva, 2010). Young leaves of plants usually have higher growth capacity and are more valuable than older ones; therefore, young leaves generally contain more defensive metabolites than older leaves (Schuman and Baldwin, 2016). Thus, insect herbivores are expected to prefer older leaves to minimize the plant resistance conferred by defensive metabolites. However, many insects still prefer and grow better on young leaves, which generally contain more nutrition, suggesting that insects have evolved strategies to minimize the negative impacts imposed by defensive metabolites (Barton and Koricheva, 2010; Kohler et al., 2015). Some studies have examined the relationship between leaf ontogeny and plant defense against the insects with chewing mouthparts, but little is known about leaf ontogeny on plant resistance to phloem-feeding insects (Barton and Koricheva, 2010; Schuman and Baldwin, 2016).

Aphids mainly feed from phloem sap of their host plants, which contains a large amount of sucrose and few essential amino acids, providing an unbalanced diet (Douglas, 2003). During their long period of coevolution, aphids have evolved strategies to cope with these constraints by increasing amino acids in the phloem and harboring symbiotic bacteria to synthesize essential amino acids (Douglas, 2003; Cao et al., 2016). Some aphids can create strong sinks at the feeding sites, increasing the flow of nutrients to the infested tissues and thereby enhancing the availability and nutrition quality of the phloem sap (Larson and Whitham, 1991; Girousse et al., 2005; Züst and Agrawal, 2016). In addition, like other feeding style insect herbivores, aphids have evolved behavioral strategies to increase their fitness by choosing particular host plants and feeding sites within them (Powell et al., 2006). Free-living aphids commonly feed on plant active sinks, like young leaves and reproductive tissues where phloem sap flows to, whereas these plant tissues generally contain more defensive metabolites (Gould et al., 2007). Therefore, there seems to be a conflict for aphids between maximizing nutrition and minimizing exposure to toxic metabolites.

Glucosinolates are the major defensive metabolites in Brassica plants, conferring resistance to most of the insects (Hopkins et al. 2009). Intact glucosinolates can be toxic to insects and their toxic effects are enhanced after hydrolysis by the enzyme myrosinase following tissue damage (Hopkins et al. 2009). Since aphids feed from phloem sap using their slender stylets, they cause tiny tissue damage to their host plants and so rarely contact these toxic breakdown products during feeding. However, indole glucosinolates have a strong antifeedant effects on the green peach aphid, *Myzus persicae* Sulzer, even without contacting myrosinase (Kim et al. 2008). Despite the negative effects of glucosinolates on aphids, *M. persicae* can grow better on plants containing higher levels of glucosinolates, suggesting that *M. persicae* may has, at least partially, adapted to these metabolites (Cole, 1997; Cao et al. 2016).

For the phloem-feeding insects, the amino acid in phloem sap is the main nutrition, and the amino acid concentration and composition as well as amino acid:sugar molar ratio in phloem sap are important indicators of nutrition quality for phloem-feeding insects (Douglas, 2003). Young leaves generally have higher phloem nutrition quality and more defensive metabolites; however, little is known about the relative importance of these two kinds of metabolites in plant resistance against aphids. We previously found that the generalist aphid, *M*. *persicae*, preferred to settle on young cabbage leaves, *Brassica oleracea* L. var. *capitata*. In order to evaluate the relative importance of nutrition and plant defense in *M. persicae-*cabbage interaction, we assessed the preference and performance of *M. persicae* on different-aged leaves of *B. oleraceae*. We also measured the glucosinolates and amino acids in different-aged cabbage leaves. Aphid feeding behavior was monitored and their honeydew excretion rate was quantified. This study intends to assess the relative importance of plant nutrition and plant defense in determining *M. persicae* feeding preference and performance on cabbage leaves.

## MATERIALS AND METHODS

### Plants and Aphids

Cabbage seeds (*Brassica oleracea* L. var. *capitata*, var. “Qingan 70”) were sown in the ground in a greenhouse (23 ± 5°C) under natural light conditions. After two months, similar-sized seedlings were transferred to 10-cm-diameter pots containing soil mixture (peat moss:perlite = 5:1) and placed in a growth chamber under a 14:10 h L/D cycle at 22 ± 2^o^C and 50% relative humidity (RH). The seedlings were watered with tap water as required and drenched with water-soluble fertilizer (2 g/L, 20-20-20 + Mg + Trace elements; COMPO Expert GmbH) every five days, receiving 5 times fertilization in total. The plants were used approximately one month after transplanting, at which time they had about 11-13 leaves. We defined “young leaves” as newly emerged leaves with a diameter smaller than 3 cm, “mature leaves” as fully expanded leaves (about the sixth to seventh emerged leaves), and “old leaves” as those closest to the soil surface. *Myzus persicae* were reared on two-month-old cabbage “Qingan 70” in cages for more than one year in the same growth chamber. If not otherwise indicated, adult *M. persicae* with similar size were used in the following experiments.

### Leaf Age and Aphid Feeding Preference

Leaf discs (1 cm diameter) were cut from different-aged cabbage leaves using a steel puncher. Two leaf discs from different age groups (i.e., young vs. mature, young vs. old, and mature vs. old) were placed in one Petri dish (9 cm diameter) lined with wet filter paper and approximately 13 apterous adult *M. persicae* were introduced to the center of these dishes. The number of aphids that had settled on each leaf disc was counted after 1, 2, 3 and 8 h.

### Leaf Age and Aphid Performance

This assay was performed with intact plants. Three apterous adult *M*. *persicae* were confined on a young, mature or old leaf using a nylon mesh bag (Cao et al., 2016). The petioles were wrapped with cotton to prevent mechanical damage by the bag. After 24 h, the adult *M. persicae* were removed, leaving five newborn nymphs on each leaf. After a further 9 d, the aphids were collected and weighed on a microbalance (resolution 0.001 mg; Sartorius MSA 3.6 P-000-DM, Gottingen, Germany), and the numbers of adults and nymphs were counted.

### Amino Acid Preferences of Aphids

*Myzus persicae* feeding preference for amino acids was assessed using 15% sucrose solution containing each of the 20 amino acids that makes up protein. Thirty-five microliter of 15% sucrose solution (control) and 15% sucrose solution containing 3 mg/mL individual amino acid were confined separately between two layers of stretched Parafilm M on a plastic Petri dish (1 cm high, 3 cm diameter). Then, 12 apterous adult *M. persicae* were introduced to each Petri dish and the number of aphids settled on each solution were recorded every 24 hours, lasting for 3 d.

### Amino Acid and Sugar Analysis

To investigate the relationship between nutrition quality and leaf age, we measured the amino acid contents in different-aged leaves and phloem sap. Amino acids in leaves were extracted by grinding the leaves in 0.05 M HCl with a glass mortar and pestle and were analyzed using an LTQ XL linear ion trap mass spectrometer (Thermo Fisher Scientific, Waltham, MA, USA) as described previously (Thiele et al., 2008; Cao et al., 2016). We used an EDTA-facilitated exudate method to collect phloem sap. The petioles of cut leaves were immersed in 800 μL of 5 mM EDTA solution (pH 7.0) for 3 h in a dark growth chamber (22°C, 100% RH). To determine the amino acid:sugar molar ratio in the phloem, we also analyzed the concentrations of sucrose, glucose, and fructose in the phloem exudate using the LTQ XL linear ion trap mass spectrometer, as previously described (Cao et al., 2016). These sugars are the main carbohydrate in the phloem exudate, while all others sugars were less than 1 mg/L and were not calculated.

### Glucosinolates Extraction and Analysis

To examine the defensive metabolite content in different-aged leaves, we analyzed the concentrations of glucosinolates in young, mature and old cabbage leaves. To inactivate the myrosinase in the leaves, 100 mg of leaves were placed in a 50-mL centrifuge tube and kept in a 96°C water bath for 3 min (Cao et al., 2016). The leaves were then ground with a glass mortar and pestle in 1 mL MilliQ water, and the mixture was centrifuged at 12,000 g, 4°C for 15 min. The supernatant was then collected and filtered through 0.22 μm syringe filters. Glucosinolates were analyzed using the LTQ XL linear ion trap mass spectrometer (Thermo Fisher Scientific, Waltham, MA, USA), as described previously (Rochfort et al., 2008; Cao et al., 2016). The relative amounts of glucosinolates were calculated according to a standard curve made by 2-propenyl glucosinolate (sinigrin). We collected aphids from different-aged cabbage leaves and analyzed the glucosinolates in their bodies as described above.

### Honeydew Excretion Assay

To investigate aphid feeding rate on different-aged leaves, we measured the dry weight of honeydew excreted by aphids feeding on young, mature and old cabbage leaves. Five adult aphids were confined to the abaxial side of each leaf using a clip cage. After 24 h, any nymphs that had been produced were removed and the clip cages were replaced by new clip cages lined with aluminum foil for a further 20 h. These clip cages were placed beneath the aphids so that honeydew that was produced dropped onto the aluminum foil. The aluminum foil was dried to a constant weight in a drying oven at 50^o^C before and after the collection of honeydew and the dry weight of honeydew produced per aphid per hour was calculated.

### Aphid Feeding Behavior

We monitored the feeding behavior of *M. persicae* on young, mature and old leaves using the Giga-8 direct-current electrical penetration graph (DC-EPG) system (W. Fred Tjallingii, Wageningen University, Netherlands). Aphid feeding activities were recorded for 8 h in a Faraday cage at 24^o^C. An 18 μm diameter gold wire was attached to the dorsum of each aphid using silver conductive glue and then aphids were placed onto leaf surface. Each adult *M. persicae* and cabbage plants were used only once. Signal was recorded by the Stylet+d software and the EPG waveforms were recognized and labeled using the Stylet+software according to Prado and Tjallingii (1994). Both software was provided by Prof. Tjallingii (Wageningen University, Netherlands). EPG parameters were calculated using the Excel workbook for automatic parameter calculation of EPG data 4.3 (Sarria et al., 2009).

### Callose Assays

Mixed instar *M. persicae* (30 mg) were confined on the abaxial side of leaves using clip cages (1 cm diameter) for 3 d. As a control, some leaves were caged without aphids for 3 d. Leaves wounded by a needle (0.2 mm diameter) for several times were positive controls. Following collection, the leaves were fixed in ethanol:glacial acetic acid (3:1) and shaken overnight (Nishimura et al., 2003). The leaves were then decolorized in 98% ethanol for 2 h and in 50% ethanol for 2 h, washed 3 times in distilled water, and stained with 0.1% (w/v) aniline blue in 75 mM phosphate buffer (pH 9.5) for 4 h in the dark. Callose deposits were viewed by a fluorescent microscope (EX 330-380 nm; DM 400 nm; BA 420 nm; Nikon eclipse 80i; Nikon Corp., Japan).

### Statistical Analysis

Aphid preferences for different-aged leaves and different artificial diets were analyzed by paired *t*-test. Levene’s and Kolmogorov-Smirnov tests were used to test homogeneity of the variances and normality of the data for aphid weight, EPG results, individual amino acid concentration in the leaves, total amino acid concentration in EDTA exudate, amino acid:sugar ratio in the phloem sap, glucosinolate concentration and honeydew weight. Any data that did not meet these tests were transformed using ln (1 = x). The effects of leaf age on these factors were then analyzed using one-way analysis of variance (ANOVA), followed by Fisher’s least significant difference (LSD) tests, at a significance level of *P* < 0.05. All statistical analyses were performed using the IBM SPSS Statistics package 19 (SPSS Inc., Chicago, IL, USA).

## RESULTS

### *Myzus persicae* Prefer and Perform Better on Young Cabbage Leaves

Significantly more aphids fed on young leaves than mature leaves after 3 h (*t* = 4.741, df = 9, *P* = 0.001) and 8 h (*t* = 5.063, df = 9, *P* = 0.001; Figure 1A), while more aphids prefer to settle on young leaves compared to old leaves since 1 h after aphid release (*t* = 3.475, df = 9, *P* = 0.007; Figure 1B). By contrast, aphids distributed evenly on mature and old leaves (Figure 1C). Compared with aphids on mature or old leaves, feeding on young leaves resulted in a higher body weight (*F* = 18.534, df = 2, 27, *P* < 0.001; Figure 1D), increased nymph production (*F* = 5.749, df = 2, 27, *P* < 0.01; Figure 1E), and an increased number of adults (*F* = 4.408, df = 2, 27, *P* < 0.05; Figure 1F) at the end of the performance assay.

**FIGURE 1.**
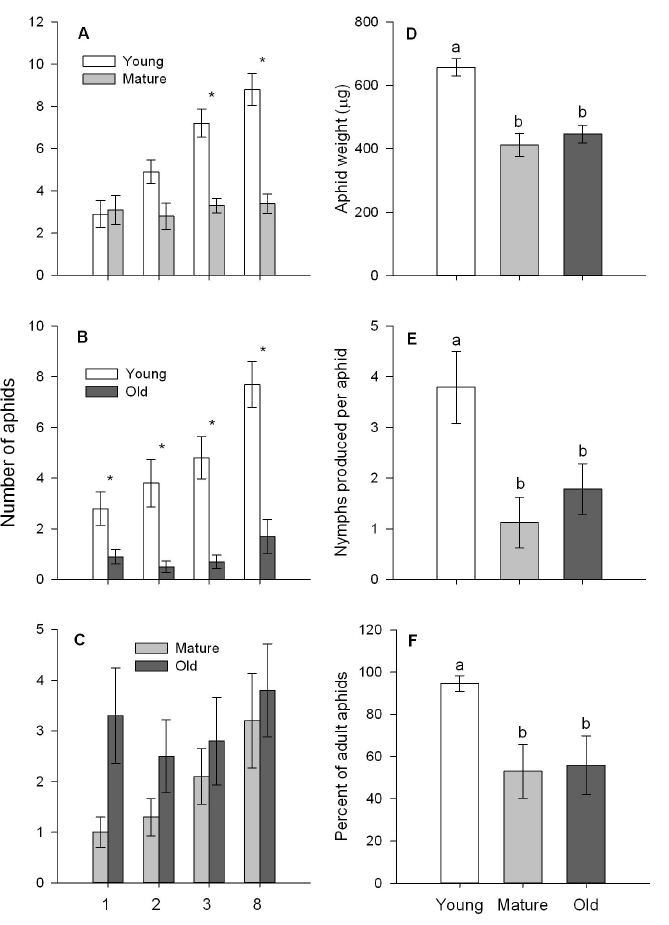
Preference and performance of *Myzus persicae* on different-aged cabbage leaves. Preference of M. persicae for pairs of different-aged leaves **(A-C)** (paired t-test: \**P* < 0.05) and performance of M. persicae in terms of body weight **(D)**, nymph production **(E)**, and number of adults **(F)**. Different letters above the bars indicate significant differences (*P* < 0.05). Values are means ± SE (n = 10).

### Young Leaves Are More Nutritious

Young cabbage leaves contained significantly higher levels of arginine (*F* = 17.569, df 2, 21, *P* < 0.001), serine (*F* = 20.829, df = 2, 21, *P* < 0.001), asparagine (*F* = 30.975, df = 2, 21, *P* < 0.001) and glutamine (*F* = 32.42, df = 2, 21, *P* < 0.001) than old or mature leaves (Figure 2A-B). The total amino acid content in the phloem exudate of old leaves was significantly lower than those of mature or old leaves, while total amino acid levels in mature and young phloem exudate were not statistically different (*F* = 12.417, df = 2, 21, *P* < 0.001; Figure 2D). Phloem sap amino acid:sugar molar ratio of young and old leaves was significantly higher than mature leaves (*F* = 15.573, df = 2, 21, *P* < 0.001; Figure 2E).

**FIGURE 2.**
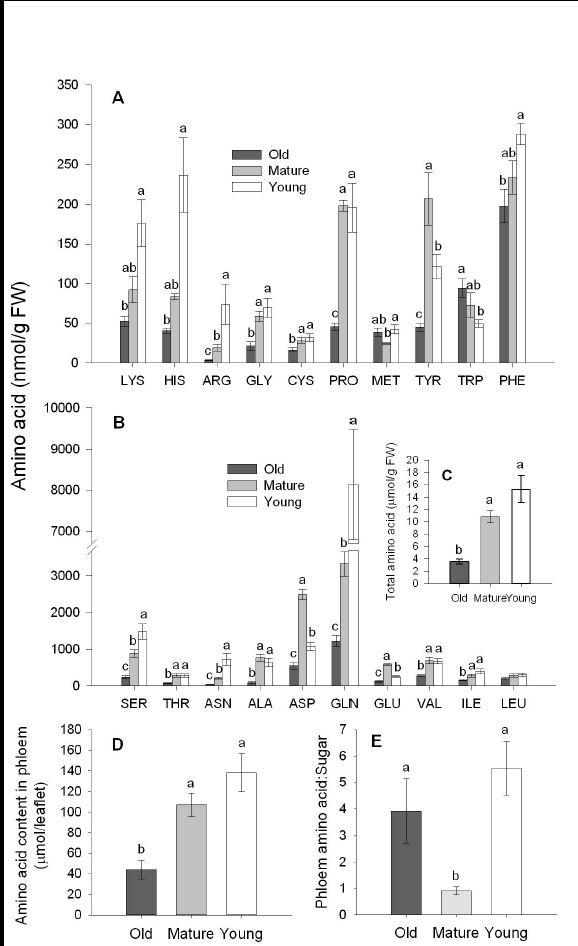
Nutrition quality in cabbage leaves for 601 *Myzus persicae*. Amino acid contents in different-aged leaves **(A-C)**, and the amino acid concentrations **(D)** and amino acid: sugar molar ratios **(E)** in the phloem sap. Different letters above the bars indicate significant differences (**P** < 0.05). Values are means ± SE (n = 8).

### Glutamine, Methionine and Valine Stimulate *M. persicae* Feeding

Significantly more aphids preferred to feed on the glutamine solution after 2 d (*t* = 2.589, df = 9, *P* < 0.05; Figure 3A) or methionine solution after 1 d (*t* = 2.632, df = 9, *P* < 0.05; Figure 3B) than the control, and the numbers of aphids chose glutamine solution or methionine solution increased with time. After 3 d, aphids also showed a significant preference for the valine solution (*t* = 2.590, df = 9, *P* < 0.05; Figure 3C
). There was no significant preference for any of the other amino acid in sucrose solution (data not shown).

**FIGURE 3.**
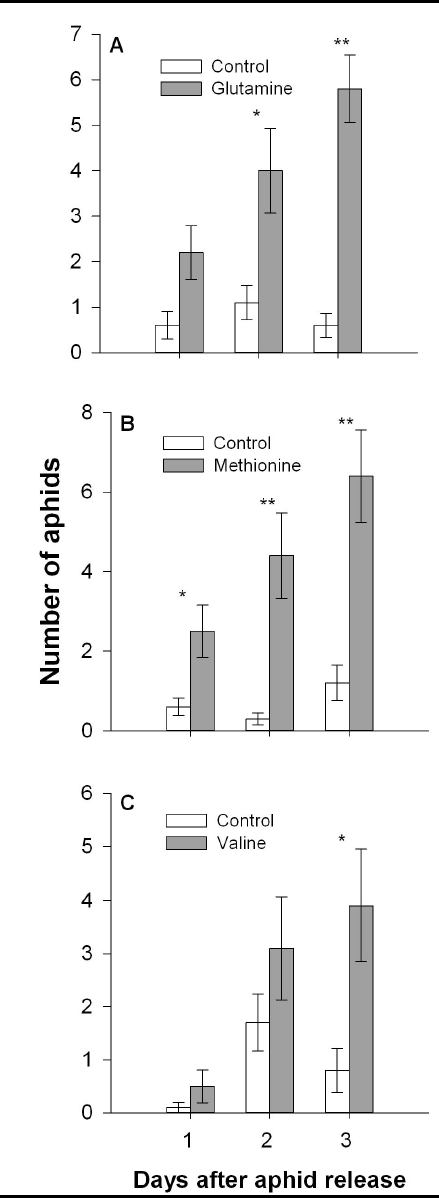
*Myzus persicae* preference for glutamine, methionine, and valine. Number of adult *M. persicae* selecting a 15% sucrose solution (control) or a sucrose solution containing glutamine **(A)**, methionine **(B)**, or valine **(C)**. (Paired t-test: \**P* < 0.05, \*\**P* < 0.01). Values are means ± SE (n = 10).

### Young Leaves and Aphids That Feed on Them Have Higher Glucosinolate Concentrations

Young cabbage leaves generally had higher glucosinolate contents than old or mature leaves, while old leaves contained the lowest levels (Figure 4A-B). Moreover, aphids that fed on young leaves had the highest glucosinolate levels in their bodies (Figure 4C-D), containing almost 2 times as much of the indole glucosinolates 1-methoxyindol-3-ylmethyl (1MI3M; *F* = 27.063, df = 2, 21, *P* < 0.001), 4-methoxyindol-3-ylmethyl (4MI3M; *F* = 53.271, df = 2, 21, *P* < 0.001) and indol-3-ylmethyl (I3M; *F* = 102.181, df = 2, 21, *P* < 0.001) than those feeding on old or mature leaves (Figure 4C-D).

**FIGURE 4.**
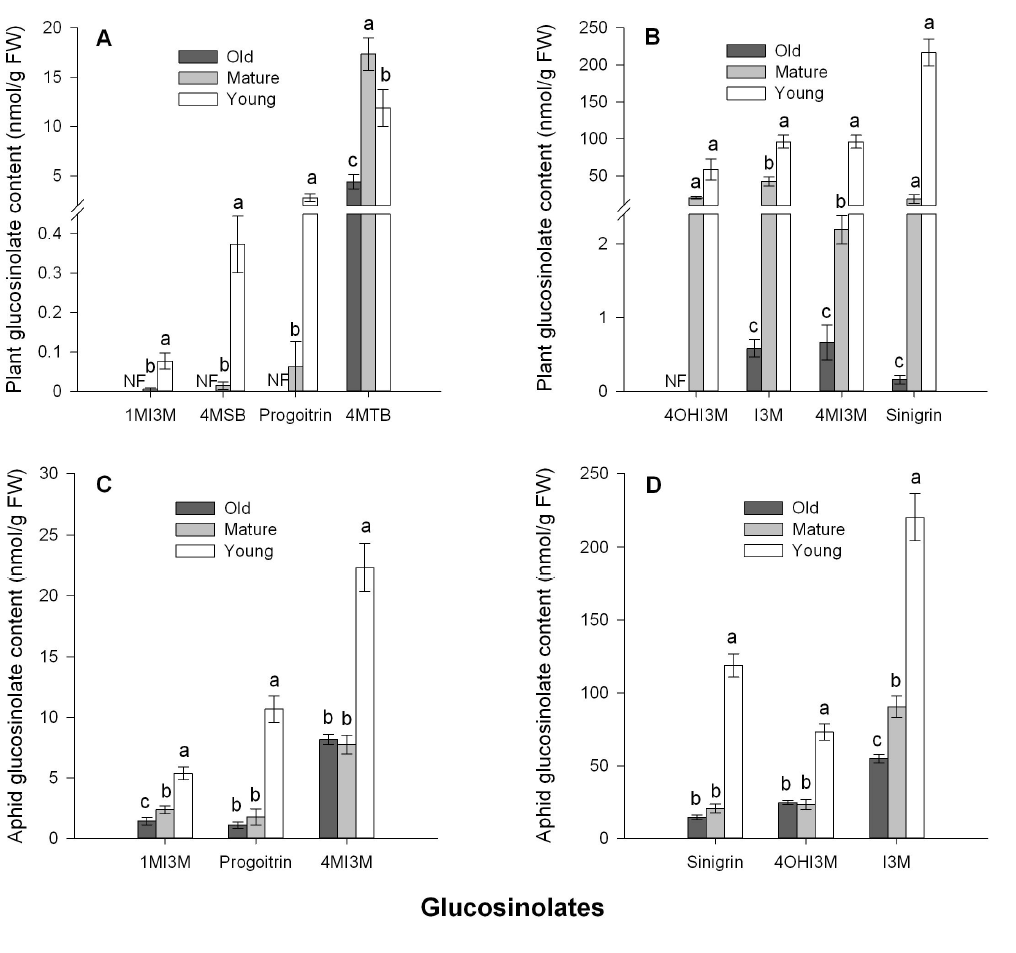
Concentrations of glucosinolates in different-aged cabbage leaves and aphids feed on them. Glucosinolates concentrations in different-aged cabbage leaves **(A-B)**, and in *Myzus persicae* feeding on these leaves **(C-D)**. Different letters above the bars indicate significant differences (*P* < 0.05). Values are means ± SE (n = 8). Glucosinolate side chain abbreviations: 4MTB, 4-methylsulfinylbutyl; I3M, indol-3-ylmethyl; 4MI3M, 4-methoxyindol-3-ylmethyl; 4OHI3M, 4-hydroxyindol-3-ylmethyl; 1MI3M, 1-methoxyindol-3-ylmethyl; 4MSB, 4-Methylsuphinylbutyl.

### *Myzus persicae* Have Longer Phloem Feeding Time and Produce More Honeydew on Young Leaves

Aphids had a significantly longer total probing time when feeding on young leaves than on old or mature leaves (*F* = 4.571, df = 2, 68, *P* < 0.05; Figure 5A). The mean phloem feeding duration of aphids on young leaves was about 4-6 times longer than those on old or mature leaves (*F* = 43.184, df = 2, 68, *P* < 0.001; Figure 5B). Aphids fed on young leaves had approximately 2 times longer total phloem feeding time than aphids on mature or old leaves (*F* = 12.638, df = 2, 68, *P* < 0.001; Figure 5C), while honeydew production rate was approximately 5 times higher from aphids feeding on young leaves than those on other leaves (*P* < 0.01; Figure 5D).

**FIGURE 5.**
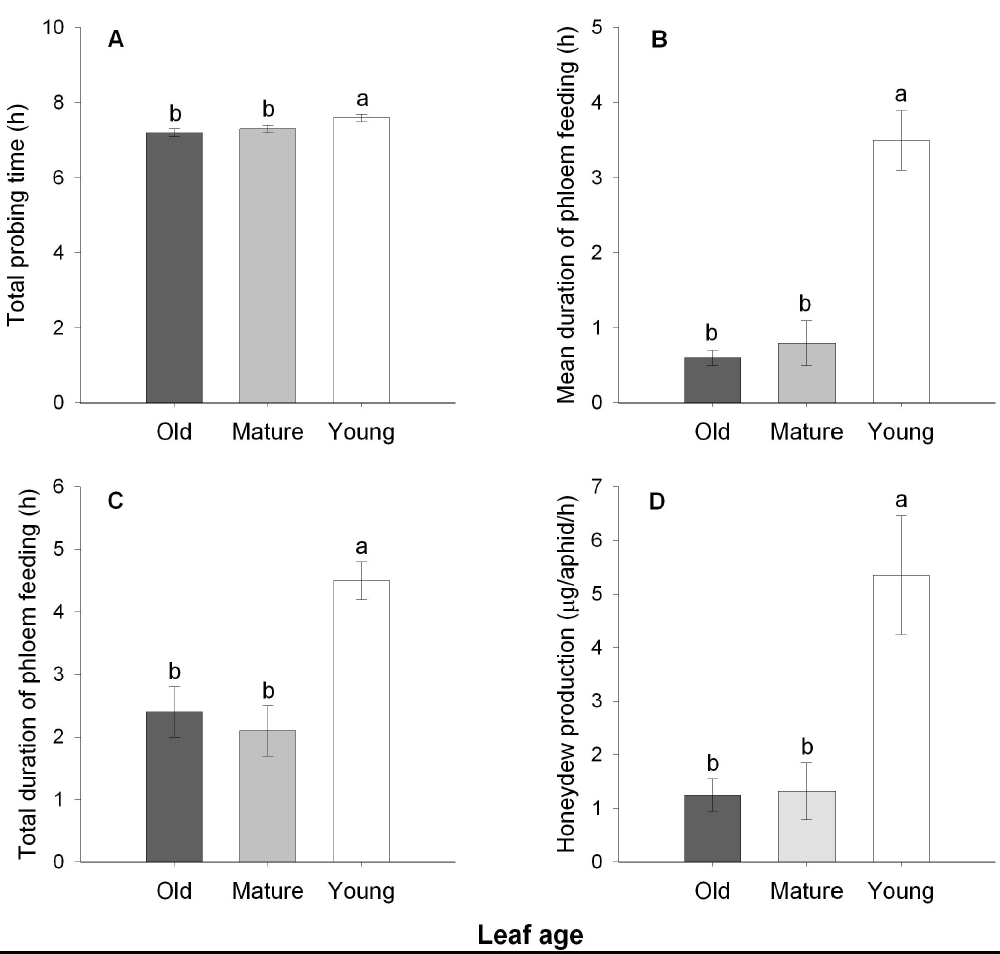
*Myzus persicae* phloem feeding activities and honeydew production rate on different-aged cabbage leaves. Total probing time **(A)**, mean phloem feeding duration **(B)**, total phloem feeding duration **(C)**, and honeydew production **(D)** by M. persicae on cabbage leaves. Different letters above the bars indicate significant differences (*P* < 0.05). Values are means ± SE (n = 22-25 for A-C; n = 5 for D).

### Aphid Feeding Behavior on Cabbage Leaves

Aphids feeding on different-aged leaves had similar numbers of probes before their stylets contacting plant phloem (*F* = 0.896, df = 2, 68, *P* =0.413; Table 1) and spent comparable time on the first probe (*F* = 0.216, df = 2, 68, *P* =0.806). Aphids had the longest phloem feeding time on young leaves, but they exhibited significantly more numbers of salivation phase (E1) (*F* = 20.914, df = 2, 68, *P* < 0.001) and phloem feeding phase (E2) (*F* = 21.162, df = 2, 68, *P* < 0.001) when feeding on old and mature leaves. In addition, *M. persicae* infestation did not elicit callose deposits on cabbage leaves (Figure S1A), while mechanical wounding resulted in callose deposits (Figure S1B).

**FIGURE 6.**
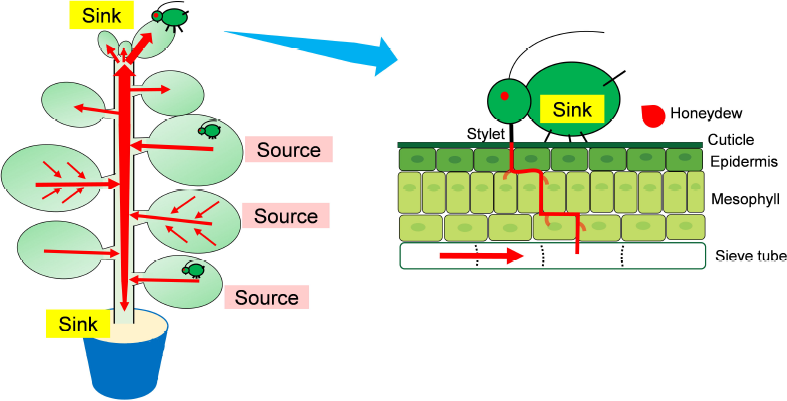
Model depicting cabbage-*Myzus persicae* interaction. Physical and chemical cues from the peripheral (nonvascular) cells control aphid preference and aphids only decide to settle or leave after penetrating the plant tissue (Powell et al., 2006). In young leaves, the higher concentration of glutamine is likely involved in host preference of *M. persicae*, while the high contents of glucosinolates act as feeding deterrents (Kim et al., 2008). Aphids are external sinks of plants and so can obtain more nutrition when feeding on young leaves, which are strong natural sinks of plants.

**TABLE 1.**
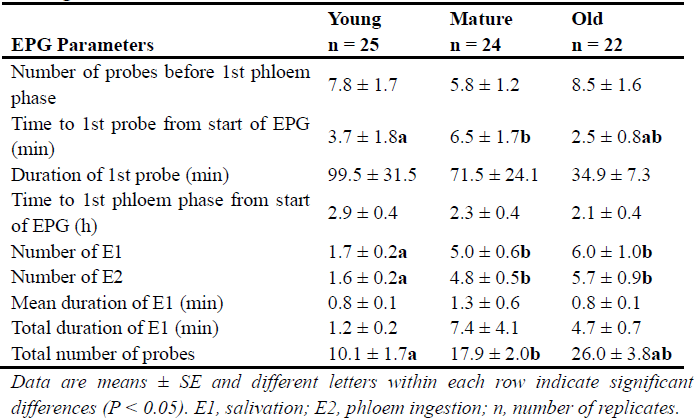
Probing behavior of *Myzus persicae* on young, mature, and old cabbage leaves.

## DISCUSSION

In this study, *M. persicae* exhibited a clear preference for young leaves, which is in agreement with previous findings for other aphid species (Gould et al., 2007; Cibils-Stewart et al., 2015). Aphids generally decide to settle on a host plant before their stylets reaching the phloem bundles (Powell et al., 2006). We found that *M. persicae* could distinguish young leaves from old leaves within 1 hour, but aphids spent at least 2 hours reaching the phloem, suggesting that cues that determine aphid preference locate at epidermal or mesophyll cells. Stimulatory and deterrent metabolites as well as plant cell wall composition are involved in host plant preference by aphids (Goggin, 2007). *Myzus persicae* showed a strong preference for the sucrose solution containing glutamine, which is present at a significant higher concentration in young leaves, suggesting that glutamine may stimulate aphid feeding and contribute to the attractiveness of young leaves for *M. persicae*. Some specialist insect herbivores rely on glucosinolates as settlement cues, which generally were present at higher levels in young leaves (Gabrys and Tjallingii, 2002; Furlong et al., 2013). However, since indole glucosinolates can act as deterrents to the generalist aphid *M. persicae*, this is unlikely to explain aphid preference for young leaves (Kim and Jander, 2007). Finally, plant cells of young leaves are at expanding stage and have flexible cell walls that may exert lower restriction to aphid stylet penetration (Divol et al., 2007).

Aphids continuously feed on plants, removing large amounts of phloem sap and reducing the hydrostatic pressure of phloem, which alters the source-sink patterns within the host plant (Girousse et al., 2005; Züst and Agrawal, 2016). According to Münch (1930), source tissues such as mature and old leaves have higher hydrostatic pressure than sink tissues such as young leaves and reproductive organs, and so it would be expected that aphids would ingest more phloem sap by feeding on source tissues. However, we found that *M. persicae* produced approximately 5 times as much honeydew per hour when they fed on young leaves, likely due to these leaves being strong natural sinks of plants that can easily draw phloem sap from source tissues during aphid feeding. The EPG results also indicated that *M. persicae* spent approximately 2 times longer feeding from phloem of young leaves than mature or old leaves. Moreover, shorter mean phloem feeding time of aphids on mature and old leaves indicated that aphids frequently withdraw their stylets from phloem of these leaves which is possibly due to reduced phloem supply caused by the depletion of phloem sap. Since *M. persicae* infestation elicits no visible callose deposits, this is not possible due to callose block of the phloem.

Amino acids are the major nutrition for aphids and are a key limiting factor for aphid growth, whose concentration and composition can influence aphid performance (Douglas, 2003). The amino acid:sugar molar ratio in the phloem sap was about 5 times higher in young leaves than in mature leaves, indicating a higher nutrition quality in phloem of young leaves. Although old leaves had a similar amino acid:sugar molar ratio in their phloem sap to young leaves, aphids obtained only one-fifth amount of phloem sap on these leaves. The higher amino acid:sugar ratio in phloem sap of old leaves may be due to the degradation of protein caused by leaf senescence (Lim et al., 2007). Reduced phloem sap ingestion is usually involved in plant resistance to aphids, suggesting the significance of phloem sap availability in determining aphid performance (Will et al., 2013). These results suggest that the amount of nutrition that aphids can obtained is as important as nutrition quality. Future research on plant-aphid interaction should pay more attention to phloem sap availability.

We found that young cabbage leaves contained the highest levels of glucosinolates, supporting the previous findings of Lambdon et al. (2003). Several studies have shown that glucosinolates are involved in plant resistance to aphids, despite aphids rarely contacting myrosinase during feeding (Kim et al., 2008; Pfalz et al., 2009). However, in our earlier study, we found that *M. persicae* infested leaves had high contents of indole glucosinolates but aphids grew better on these leaves (Cao et al., 2016). Similarly, the cabbage aphid, *Brevicoryne brassicae* L., also performs better when feeding on reproductive tissues (flowering canopy) than on vegetative tissues (leaves), despite the flowering canopy having higher levels of glucosinolates (Cibils-Stewart et al., 2015). and in this study, *M. persicae* grew better on young leaves that contained higher levels of glucosinolates. These findings suggest that the effects of glucosinolates in plant resistance to aphids has significant variation, which may be due to nutrition compensate or aphid adaption. And aphids tend to feed on more nutritious tissues to maximize their fitness, regardless of the higher levels of toxics in these tissues. In addition, aphids generally had higher levels of I3M and 1MI3M in their bodies than leaves they were feeding on, but aphids feeding on young leaves contained significantly lower 4MI3M, a more toxic indole glucosinolates for *M. persicae*, than that in young leaves, implying that aphids may have converted this glucosinolates to lesser toxic metabolites.

Both toxic metabolites and nutrition are essential factors in plant resistance to aphids, but aphids have evolved strategies to cope with these constraints by detoxifying the toxics and selecting feeding sites (Dreyer and Campbell, 1987; Mathers et al., 2016). We found that young cabbage leaves contained higher levels of glucosinolates, but also provide more nutrition, contributing to the improved performance of *M. persicae*. The source-sink relationship within plants influences phloem sap availability and thus aphid performance, which deserves more attention in the research of aphid-plant interaction.

## AUTHOR CONTRIBUTIONS

HC originally formulated the idea and designed the experiments. HC and ZZ performed the experiments. HC did the statistical analyses and wrote the manuscript. XW and TL provided editorial advice. TL provided funding for the project.

## ACKNOWLEDGMENTS

We are grateful for the assistance of all the members in the Key Laboratory of Applied Entomology, Northwest A&F University, Yangling, Shaanxi, China.

